# Linking continuous behavior to aesthetic enjoyment in a walkable virtual-reality museum tour: effects of agency and a painting-level analysis framework

**DOI:** 10.64898/2026.08.27.747505

**Authors:** Yana Sklyar, Joseph Hendler, Tom Schonberg

## Abstract

Museum visits typically follow curator-defined routes that constrain how visitors shape their own experience, yet choice is widely held to heighten engagement, autonomy, and enjoyment. Virtual reality (VR) offers a setting in which to study these processes because it combines ecological immersion with precise, continuous behavioral measurement. We investigated (i) whether VR-derived behavioral signals are associated with self-reported enjoyment during a virtual museum tour, and (ii) whether the level of agency afforded to visitors influences enjoyment. Forty-eight adults completed a room-scale, life-size VR tour (8 × 4 m) of seven paintings from the Tel Aviv Museum of Art, each accompanied by a synchronized audio guide. Synchronized gaze and head-position streams were logged continuously (50 Hz) and segmented into painting-level viewing episodes using a trial-and-tile pipeline that intersects each painting’s trial interval with an empirically defined spatial window in front of the canvas. Participants were randomly assigned to one of three agency conditions, Active (choice before every artwork), Semi-Active (choice for the first three), or Passive (fixed route),while the artwork sequence was held identical. Self-reported enjoyment at the tour and painting levels did not differ reliably across agency conditions. Among VR-derived measures, gaze engagement during the audio guide showed the clearest (though modest) association with painting-level liking, whereas locomotion and pacing measures were weak and inconsistent predictors. Agency nonetheless reliably modulated several gaze-and time-based viewing measures. The findings reveal a dissociation between subjective enjoyment and the micro-structure of viewing, and establish a reusable framework for full-tour, painting-level behavioral analysis in immersive settings.

## 1 Introduction

Digital access to museum collections has expanded rapidly over the past decade, and platforms such as Google Arts & Culture have offered virtual walks through real galleries since 2011 [1]. The COVID-19 pandemic accelerated this trend, prompting many institutions to expand virtual tours and digital experiences to keep audiences connected to their collections [2]. In most web-based formats, however, visitors “go to the museum” through a screen, navigating by teleportation or game-like controls [3]. Such systems greatly increase access but provide a limited sense of bodily presence and only a narrow window onto how people actually move and look [4, 5]. This raises a methodological question: if museum visits increasingly move into digital formats, what tools allow viewing behavior and experience to be studied with greater ecological validity and richer measurement than a standard screen-based tour can offer?

VR has become an attractive platform for studying art perception and cultural heritage in controlled, instrumented environments. Gaze patterns recorded in a virtual replica of an installation artwork can closely resemble those obtained in the physical installation [6], and dual-mode virtual museums have been evaluated for navigation efficiency, usability, and satisfaction [7]. A subsequent line of work moved from desktop and mobile experiences to head-mounted displays (HMDs) to test how immersion changes the visit. Immersive VR can enhance presence, engagement, and motivation relative to 360° video and mixed-reality formats [5]; immersive HMD environments support more accurate cognitive maps and higher reported aesthetic experience than desktop versions [8]; and wearable HMDs outperform smartphone-based VR on immersion and downstream attitudes [9]. Across these studies, locomotion is typically mediated by controllers or teleportation, and outcomes are measured mainly at the tour level (presence, learning, user experience).

In consumer and retail research, immersive VR stores with body tracking yield stronger telepresence and more natural interaction than desktop versions [10]; comparing physical walking to teleportation, locomotion technique left emotional states and purchasing outcomes largely unchanged despite clear differences in movement [11]; and walkable VR supermarkets reproduce real-world food-selection patterns with only minor differences in information seeking [12]. In the art domain, free-walk trajectories around a single abstract artwork can be modeled with deep-learning methods to anticipate movement [13], and combining free walking with head-, hand-, and eye-tracking reveals distinctive approach and viewing patterns around a single VR painting [14]. Together, these studies show that immersive VR can link movement to choice and preference, but they also reveal a gap: real-walk measurement has rarely been embedded in a full museum-like tour where behavior unfolds across multiple artworks and can be related both to the visit as a whole and to preferences for individual works.

Building on this work, we extend real-walk VR from single-artwork encounters to a full virtual museum tour with a synchronized audio guide and continuous multimodal logging. We implemented a walkable gallery containing seven life-size paintings and combined natural walking with continuous gaze logging and trial-based segmentation, producing a multimodal pipeline that links in-tour behavior to both overall visit satisfaction and painting-level enjoyment. We additionally introduced an experimental manipulation of agency. Because choice tends to enhance intrinsic motivation and enjoyment [15–20], participants were randomly assigned to Active, Semi-Active, or Passive conditions; critically, the artwork sequence was identical across conditions, so the manipulation altered the experience of choosing rather than the content of the tour.

We addressed two research questions. First, are VR-derived behavioral signals associated with self-reported tour enjoyment and with single-image preferences? We expected that signals capturing how participants looked at and moved around artworks would show systematic relationships with tour satisfaction and painting-specific liking. Second, does the level of agency affect self-reported enjoyment? We hypothesized that higher agency (Active > Semi-Active > Passive) would increase tour satisfaction and related attitudes (willingness to revisit, to recommend, and to seek similar experiences).

## 2 Methods

### 2.1 Participants

We recruited adults (18–39 years; all genders) from Tel Aviv University and the surrounding community, with no history of neurological or psychiatric disorders. All participants provided written informed consent and received monetary compensation (40 NIS per hour). The experiment was approved by the Tel Aviv University ethics committee. Data were collected from 67 participants; after quality-control exclusions (Sect. 2.8) the final sample comprised 48 complete datasets (N = 48; 336 painting episodes).

### 2.2 Virtual environment and apparatus

The virtual museum ran on a Meta Quest Pro HMD, which tracks three-dimensional head position and orientation and supports built-in eye tracking. The application was developed in Unity (2022.3.18f) using licensed Unity Asset Store packages customized in Blender (3.6.8 LTS); gallery wall geometry was rescaled in Blender to match the physical room. The walkable area was 8 × 4 m, performed in the TAU XR Arena, enabling natural, room-scale locomotion. The tour comprised seven works from the museum’s Mizne–Blumental collection (Table 1), presented at their real physical size. Artwork textures provided by the museum were upscaled for in-headset fidelity using MidJourney (v6, resolution enhancement only; no semantic edits). Each artwork had a synchronized audio-guide track recorded by co-author J. Hendler (durations 48.6–87.3 s). All on-screen and in-headset prompts were presented in Hebrew.

**Table 1.**
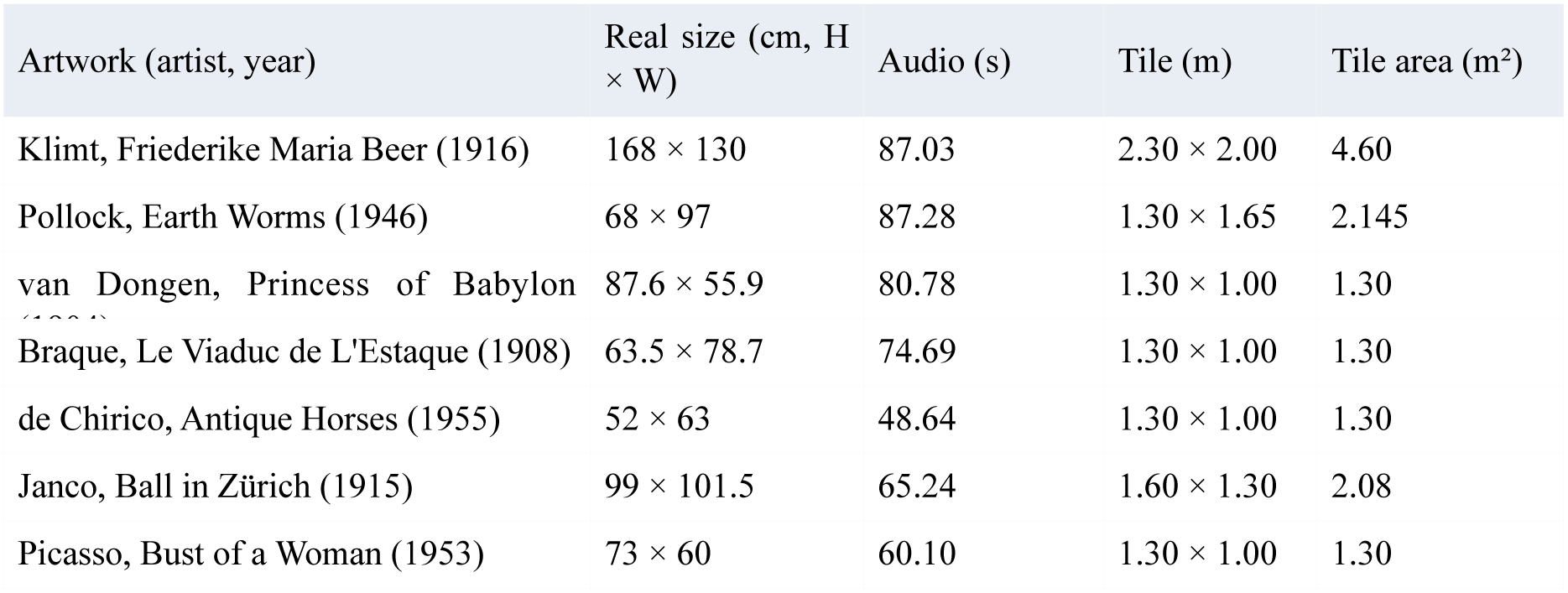
Artwork stimuli (fixed presentation order) and the empirically defined spatial “tiles” used for painting-level segmentation.

### 2.3 Agency manipulation

Participants were randomly assigned to one of three between-subjects conditions. In the Active condition, after each artwork participants answered an in-headset *“*choice” question about a feature of the next work they wished to see (three options per screen; Fig. 1, Fig. 2). In the Semi-Active condition, choice questions appeared for the first three artworks, after which the route became passive. In the Passive condition, the curated route was fully pre-set with no choice questions. Because every option led to the same next painting, the artwork sequence (Klimt, Pollock, van Dongen, Braque, de Chirico, Janco, Picasso) was identical across conditions. This design isolates the experience of choosing while holding content exposure constant and provides variation in exploration patterns for the methodological analyses.

**Fig. 1.**
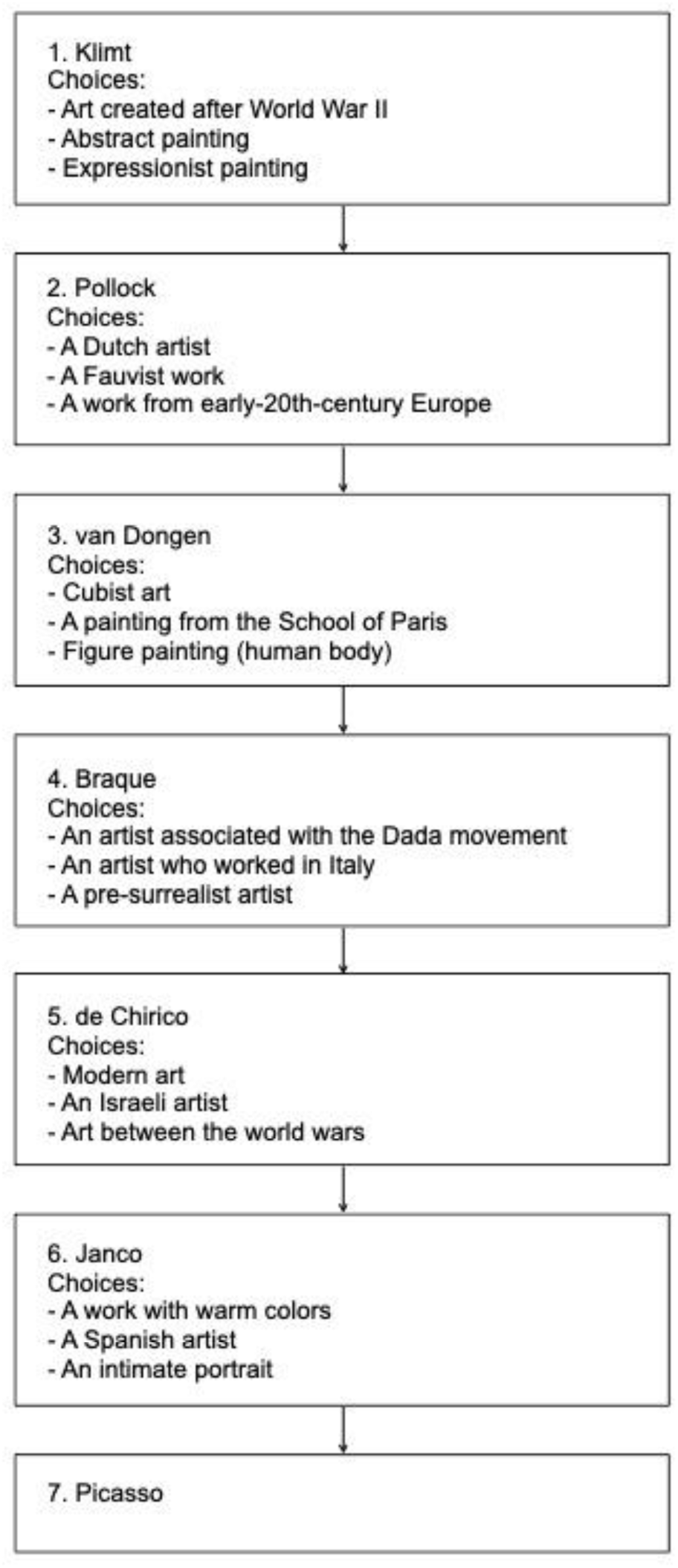
Choice-question flow across the seven artworks (Active and early Semi-Active conditions). Each box lists the feature options offered before the next painting; every option led to the same next work, so the sequence was identical across conditions.

**Fig. 2.**
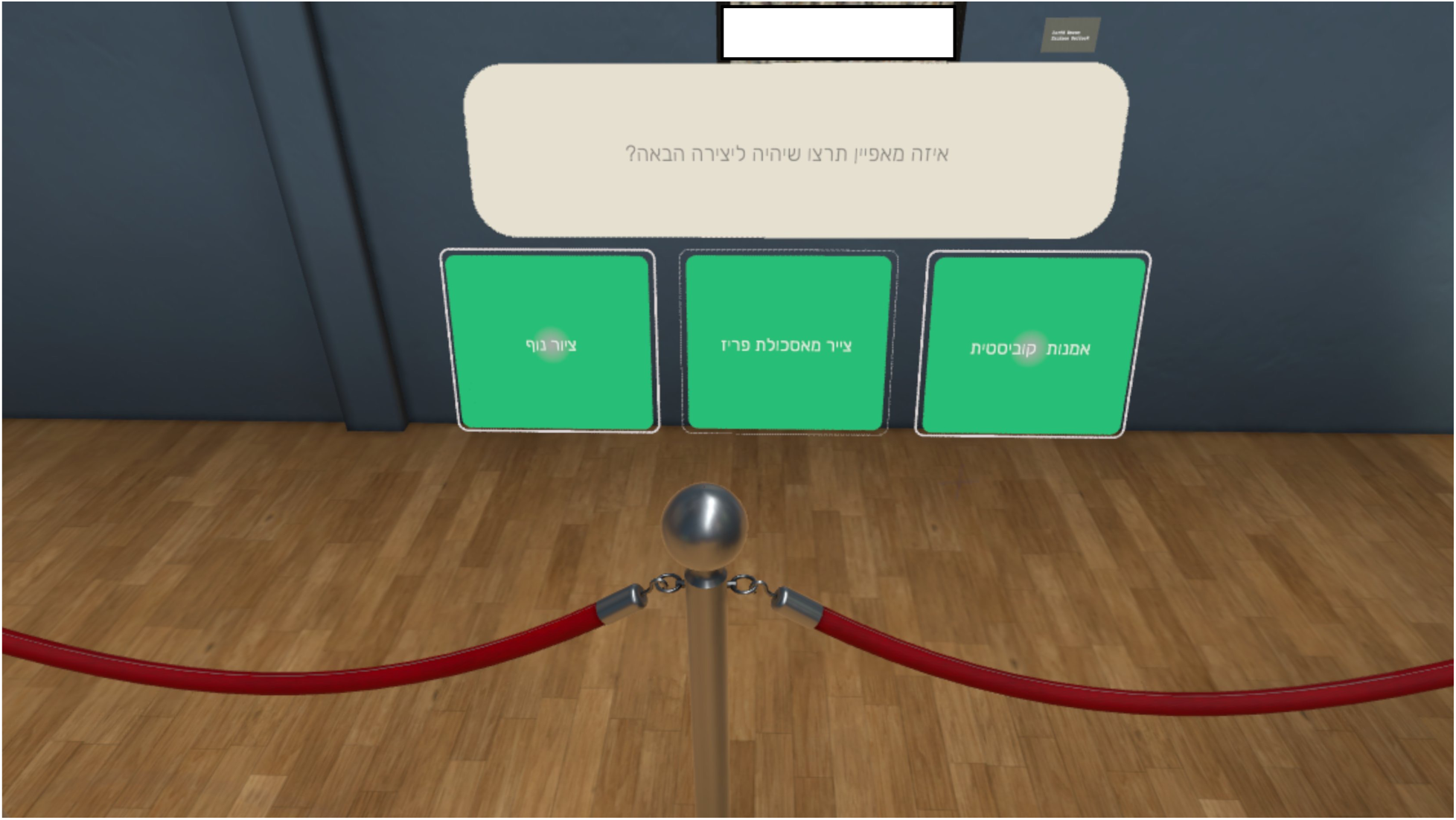
Example in-headset choice screen shown during the tour. The Hebrew prompt asks which feature the participant would like the next artwork to have, with three option buttons below the current painting.

### 2.4 Procedure

After consent, participants completed a Hebrew pre-tour questionnaire on demographics and museum habits. They donned the headset, completed the built-in Meta eye-tracking calibration, and performed a brief navigation demo in the same room with three empty frames in place of artworks. A recorded voice (the audio-guide voice) instructed participants to follow a green arrow to each frame, press a wall-mounted button to trigger an explanation, and practice answering a choice question and pressing virtual buttons. After the second frame the demo ended, the empty frames disappeared, the real paintings appeared in the same locations, and the tour began. A minor mismatch between a virtual button and its collider at the start sometimes required participants to reach slightly *“*deeper” into the wall; because this occurred during the demo, participants adapted quickly and no further button-press difficulties were observed during the main phase.

After the tour, participants completed a Hebrew post-tour questionnaire on a desktop computer (Google Forms). The first part assessed the tour as a whole (overall satisfaction, satisfaction with the explanations, perceived duration, willingness to hear more explanations, to revisit, to visit other museums with similar guidance, and to recommend the experience), all on 7-point scales. The second part presented each of the seven artworks with its image for a 1–7 liking rating. The final part was a binary choice preference task: the first seven pairs each presented a tour painting alongside an unseen painting by the same artist (an exposure/validity check), and the remaining pairs presented only unseen paintings (collected for future methodological use). Sessions lasted approximately 20 min; VR logs and questionnaires were exported after each session.

### 2.5 Signal logging and synchronization

All in-headset streams were exported to CSV files sharing a common time base (public templates at minervaxr.sites.tau.ac.il/open-science); each file’s first time column was converted to numeric seconds from scene onset. The single time axis aligned the continuous head/gaze stream (ContinuousData, sampled at 50 Hz), audio-guide events (AudioGuideTiming), in-experience logs (e.g., instruction screens), and in-headset questions and answers, so that gaze and movement could be matched to the same moments in the tour. Processed tables and the full analysis code are available in the project repository (github.com/SlabMuseum/Enjoyment).

### 2.6 Trial-and-tile segmentation

For each participant, the experiment was parsed into seven trials, one per artwork, from headset logs, audio-guide timing, and interaction events. Trial boundaries were defined relative to the shared time axis and differed slightly by condition (Fig. 3). For all conditions, the first trial (Klimt) began when the *“*let’s start the tour” board was hidden. In the Active condition, each subsequent trial began immediately after the participant chose the next artwork and ended when that painting’s audio guide finished. In the Semi-Active condition, the first artworks followed the Active pattern, after which trials followed a passive pattern (starting at the end of the previous audio guide). In the Passive condition, all trials began at tour onset or at the end of the previous audio guide and ended when the current painting’s audio guide finished. By construction, the painting-level trial windows exclude the choice screens while preserving the core viewing episode (walking to the artwork, exploring it, and listening to the audio guide).

**Fig. 3.**
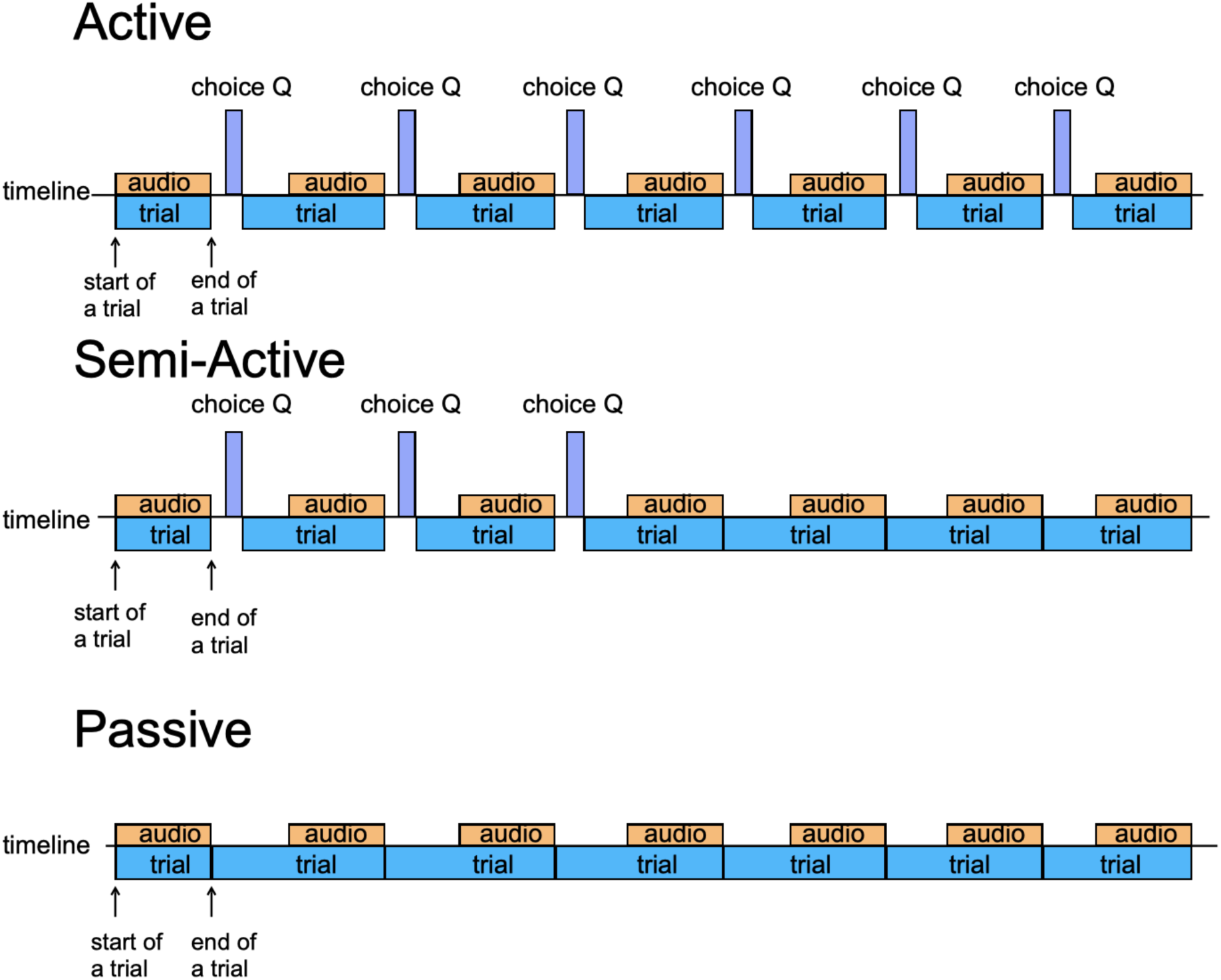
Schematic trial segmentation per condition (Active, Semi-Active, Passive), showing the trial, audio-guide, and choice-question intervals on a shared timeline. Painting-level trial windows exclude the choice screens.

To connect behavior to specific artworks, participant trajectories were exported and visualized; these revealed a consistent *“*footprint” in front of each painting where participants tended to stand while viewing and listening (Fig. 4). Square tiles were defined around each painting from these observed standing regions, larger tiles for the two large-scale works (Klimt, Pollock) and smaller tiles for the rest (Table 1). A moment was labeled *“*in tile” when the head position (X, Z) fell within the stored tile bounds. Final painting-level segments were the intersection of (i) the painting’s trial interval and (ii) the in-tile timestamps, and were used to compute gaze, movement, and (for future work) facial-expression measures per artwork.

**Fig. 4.**
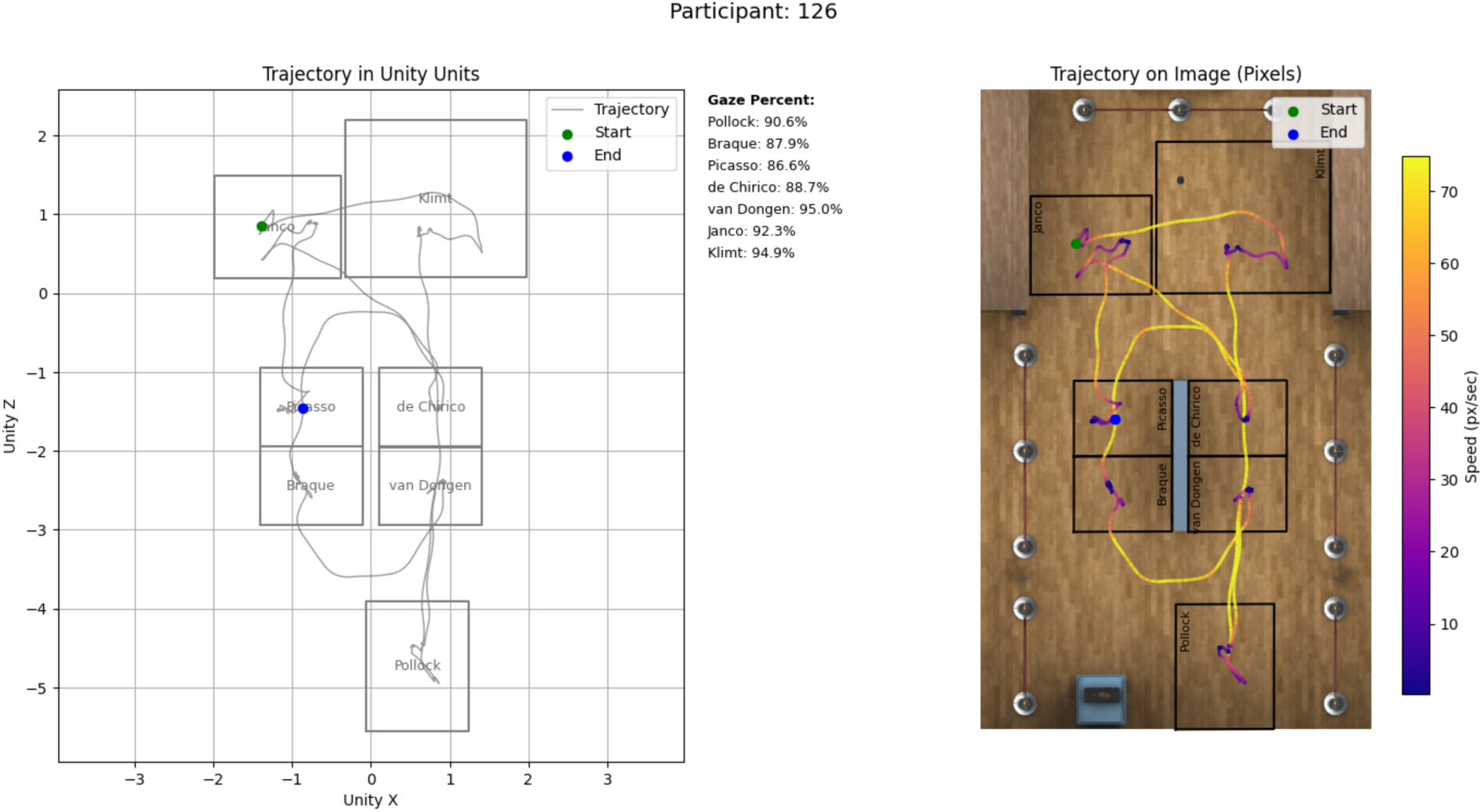
Example participant trajectory in Unity coordinates (left) and overlaid on the gallery floor plan (right), colored by speed, with per-painting gaze percentages. These footprints motivated the empirically defined tiles.

### 2.7 Behavioral measures

Gaze was resolved by ray casting: at each sample, Unity recorded which collider the gaze ray hit (FocusedObject). Because invisible demo colliders remained near the real artworks, gaze that visually landed on a real canvas was sometimes attributed to a demo object; we corrected this using the hit position and known collider bounds, relabeling such hits to the correct artwork (CorrectedFocusedObject). From the segmented windows we derived: GazeTime (cumulative on-canvas gaze time, computed by multiplying the count of on-painting samples by the 0.02 s sampling interval); GazePercent_Audio (percentage of the audio-guide interval with gaze on the painting); SaccadeRate, a proxy for visual scanning [21, 22] (the number of gaze-position shifts exceeding 0.01 m between consecutive 3-D gaze-hit samples, divided by audio-guide duration, following threshold-based event-detection approaches that separate true gaze shifts from measurement noise [23]; the 0.01 m threshold corresponds to roughly 0.3–0.6° of visual angle at typical viewing distances of ∼1–2 m, comparable to the spatial uncertainty of HMD eye tracking [23, 24]); and ReactionTime (latency from segment start to the first sustained fixation on the canvas). Saccade rate was computed during the audio-guide segment because the trial window also includes navigation, during which a participant may already be in the tile but not yet looking at the artwork. For interpretation only, saccade rates can be grouped into rough engagement bands, with sustained attention typically accompanied by fewer saccades [25]; all inferential tests used the continuous values.

Spatial measures were TimeAtTheTile (duration of the painting’s viewing episode) and TileTimePercent_Audio (TimeAtTheTile as a percentage of the audio-guide duration). Locomotion measures were computed from head position across the full session: instantaneous walking speed (ground-plane displacement divided by inter-sample interval), total distance, average speed, and time still versus moving, using a 0.005 m/s stillness threshold to suppress tracking jitter, consistent with threshold-based movement detection in VR sensor streams [26]. Because the paradigm relied on natural room-scale walking, locomotion summaries are interpreted in light of known variability in natural-and redirected-walking VR [27]. Participant-level aggregates were means or sums across the seven paintings (e.g., AvgGazeTime, AvgGazePercent_Audio, AvgSaccadeRate, TotalDistance), together with an AvgGeneralRating index averaging the seven general post-tour items.

### 2.8 Quality control and statistical analysis

Sessions were excluded for corrupted ContinuousData logs that prevented valid trajectory reconstruction (3), missing or incomplete eye-tracking calibration (12), or an application crash mid-tour (4). Extracted trials were validated to ensure StartTime preceded EndTime, and trajectory plots were inspected to confirm data plausibility before inclusion. Occasional *“*teleportation” jumps in recorded head position were addressed in-session by pausing, re-calibrating the play space, and resuming; because these did not affect timestamps, tiles, or gaze labelling within trials, the affected sessions were retained, but gross locomotion metrics (total distance, average speed, total duration) are interpreted with caution. The headset also recorded facial-muscle signals; these were not analyzed here but are time-aligned to the same axis for future segmentation.

Analyses were performed in Python 3.12.0 (pandas 2.3.1; SciPy 1.13.1 for non-parametric tests and correlations; scikit-posthocs 0.11.4 for post-hoc comparisons; Matplotlib 3.10.5 and Seaborn 0.13.2 for figures). Effects of tour type were tested with Kruskal–Wallis tests across Active, Semi-Active, and Passive conditions. Associations between enjoyment and VR-derived variables were assessed with Spearman correlations at both the painting level (painting liking vs. painting-level features) and the participant level (average general rating vs. aggregates).

## 3 Results

### 3.1 Descriptive overview

Because each valid participant contributed one segment per painting, the painting-level dataset was complete and balanced (N = 336 segments; 48 participants × 7 paintings). The segmentation produced sensible distributions with substantial variability across both subjective and behavioral measures (Table 2). Gaze during the audio guide was typically high (median Gaze Percent_Audio 81 %) while still allowing large individual deviations. TileTimePercent_Audio frequently exceeded 100% (median 116%), indicating that many participants remained in front of the painting after the narration ended rather than moving on immediately; this is expected by design, since the tile window also captures time before and after the audio guide. At the participant level (N = 48), the mean tour lasted about 10.8 min (TotalExperimentTime mean 649 s), and participants differed markedly in how much they moved (total distance median 61 m, range ∼36–138 m), confirming meaningful variation in exploration style rather than a single stereotyped route.

**Table 2.**
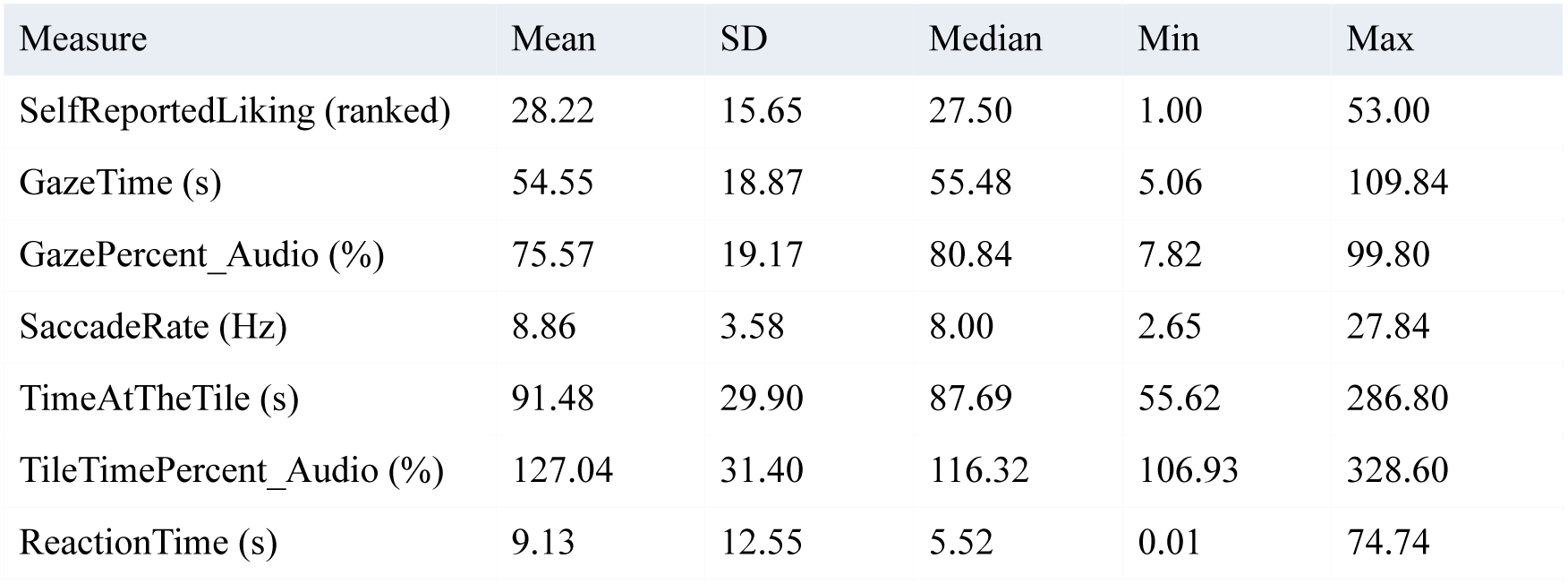
Descriptive statistics for painting-level segments, pooled across paintings and conditions (N = 336).

Reaction time to first fixation varied strongly across paintings, with longer, more variable values for Klimt than for Janco or Picasso, suggesting that some artworks elicited slower settling into the canvas at the start of the episode (Fig. 5). Plotting GazeTime against each painting’s audio-guide duration showed that audio lengths set an upper bound on guided looking and that, even within the audio window, looking was not continuous: for every painting, gaze time fell below the narration duration for many participants (Fig. 6). This is precisely the moment-to-moment variability a VR pipeline should capture and confirms that the measures reflect realistic viewing behavior rather than assuming *“*listening = looking.”

**Fig. 5.**
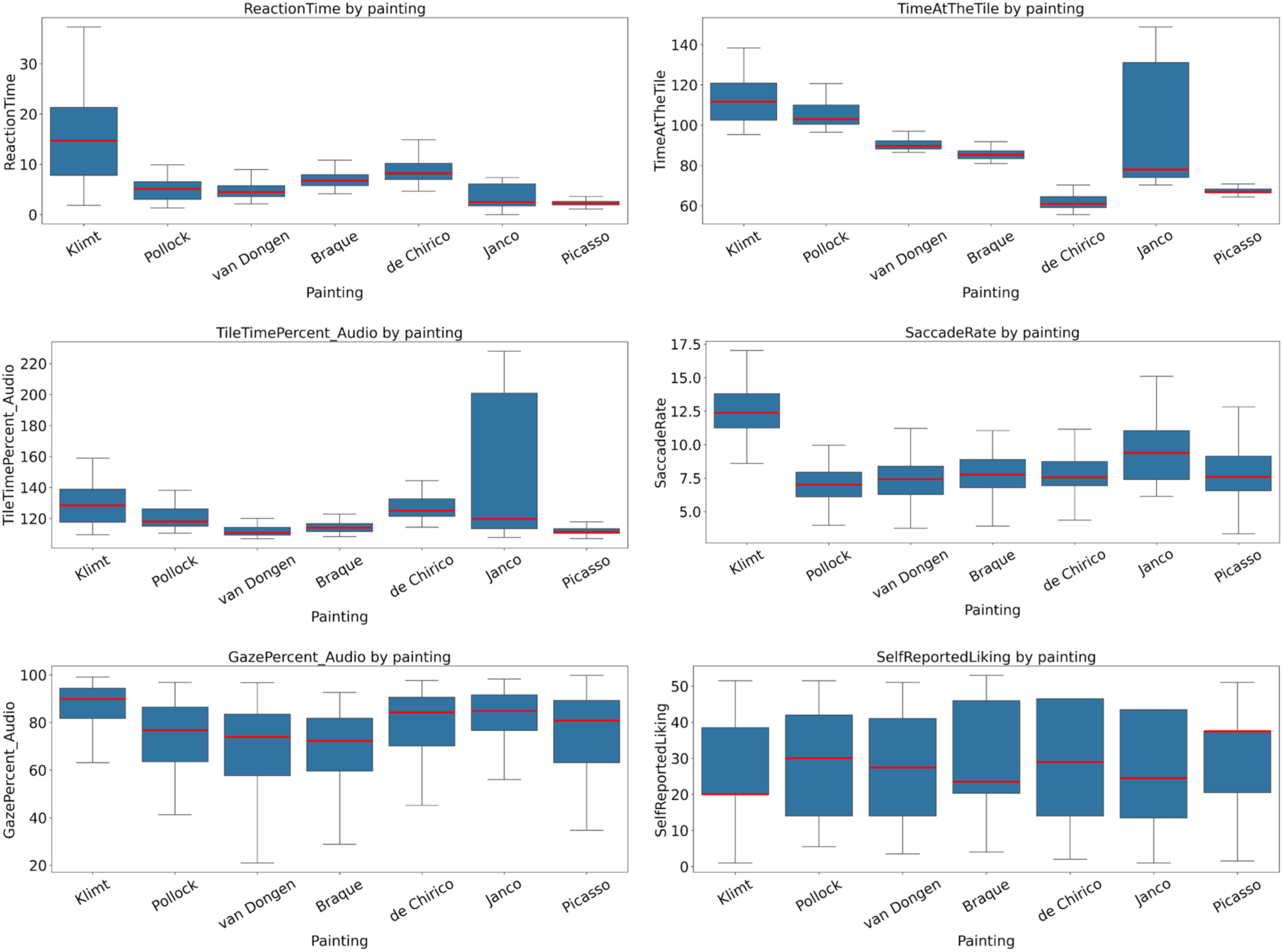
Descriptive distributions of painting-level measures across the seven artworks (medians in red): reaction time, time at the tile, in-tile time as a percentage of audio duration, saccade rate, gaze during the audio guide, and self-reported liking.

**Fig. 6.**
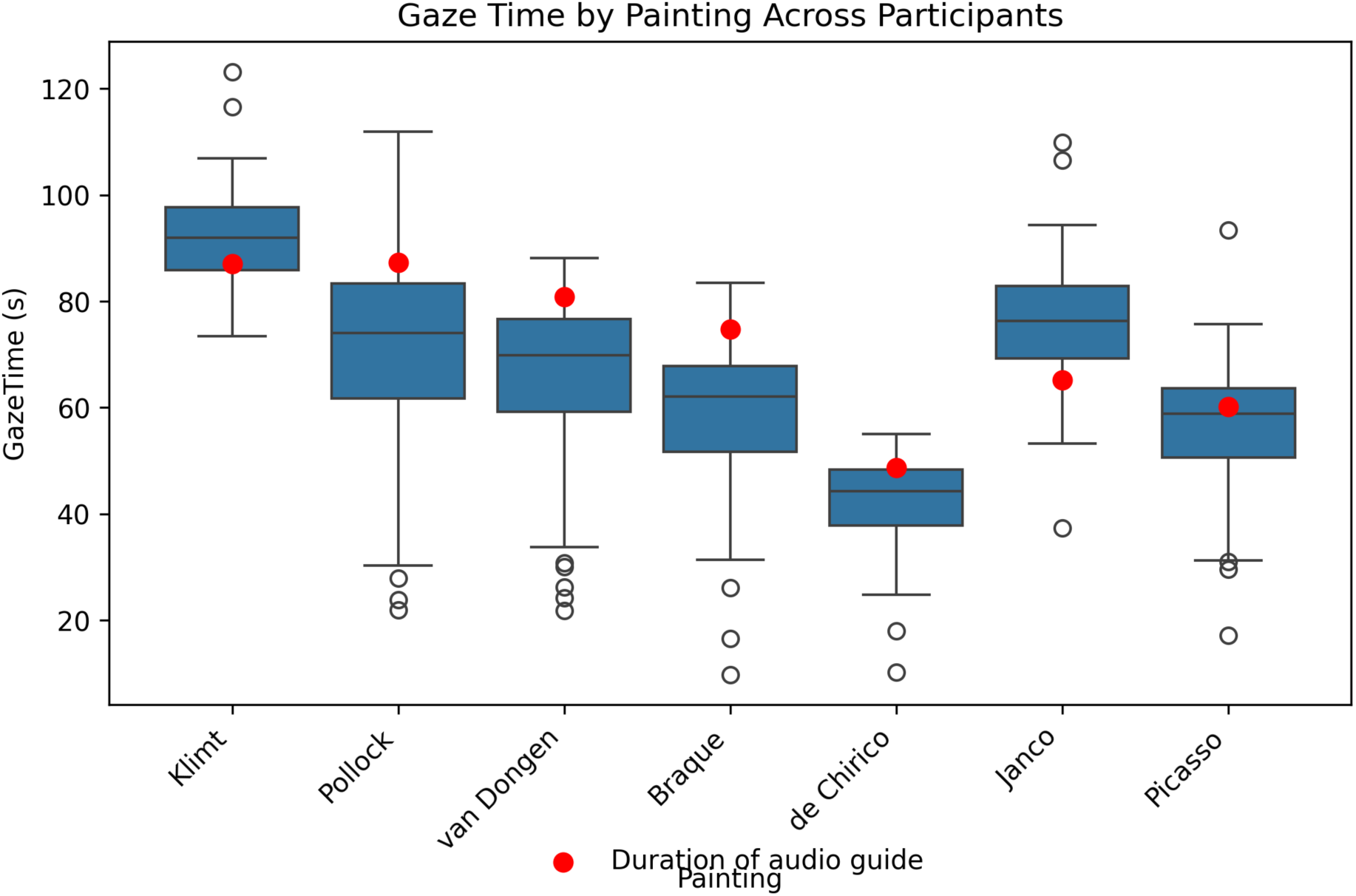
Gaze time by painting across participants, with each painting’s audio-guide duration overlaid (red points). For every painting, gaze time falls below the narration duration for many participants, indicating that looking is not continuous during the audio guide.

**Fig. 7.**
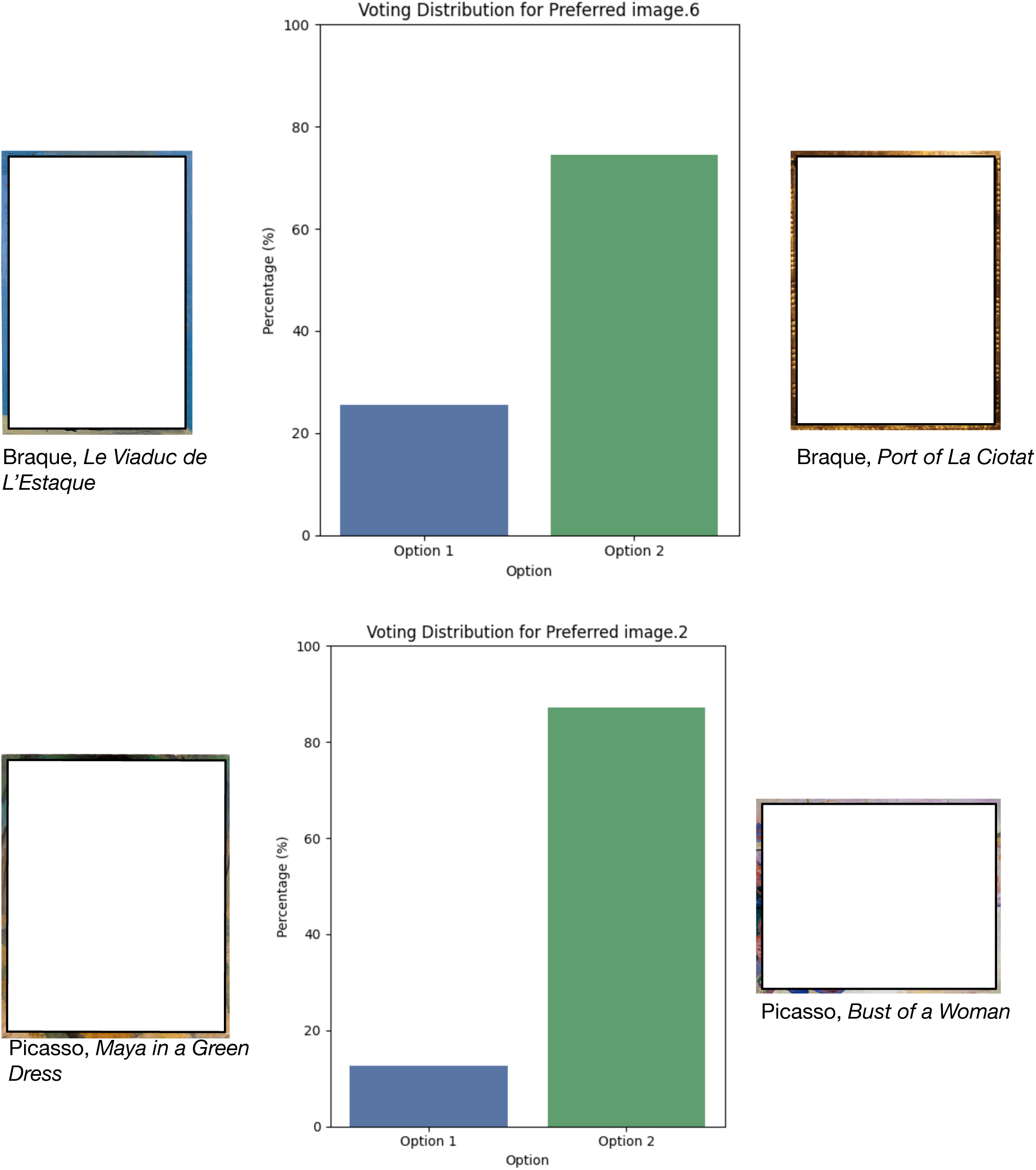
Binary choice task: voting distributions for tour paintings paired with an unseen work by the same artist. The proportion choosing the tour painting ranged from 0.13 (Braque) to 0.75 (Picasso).

**Fig. 8.**
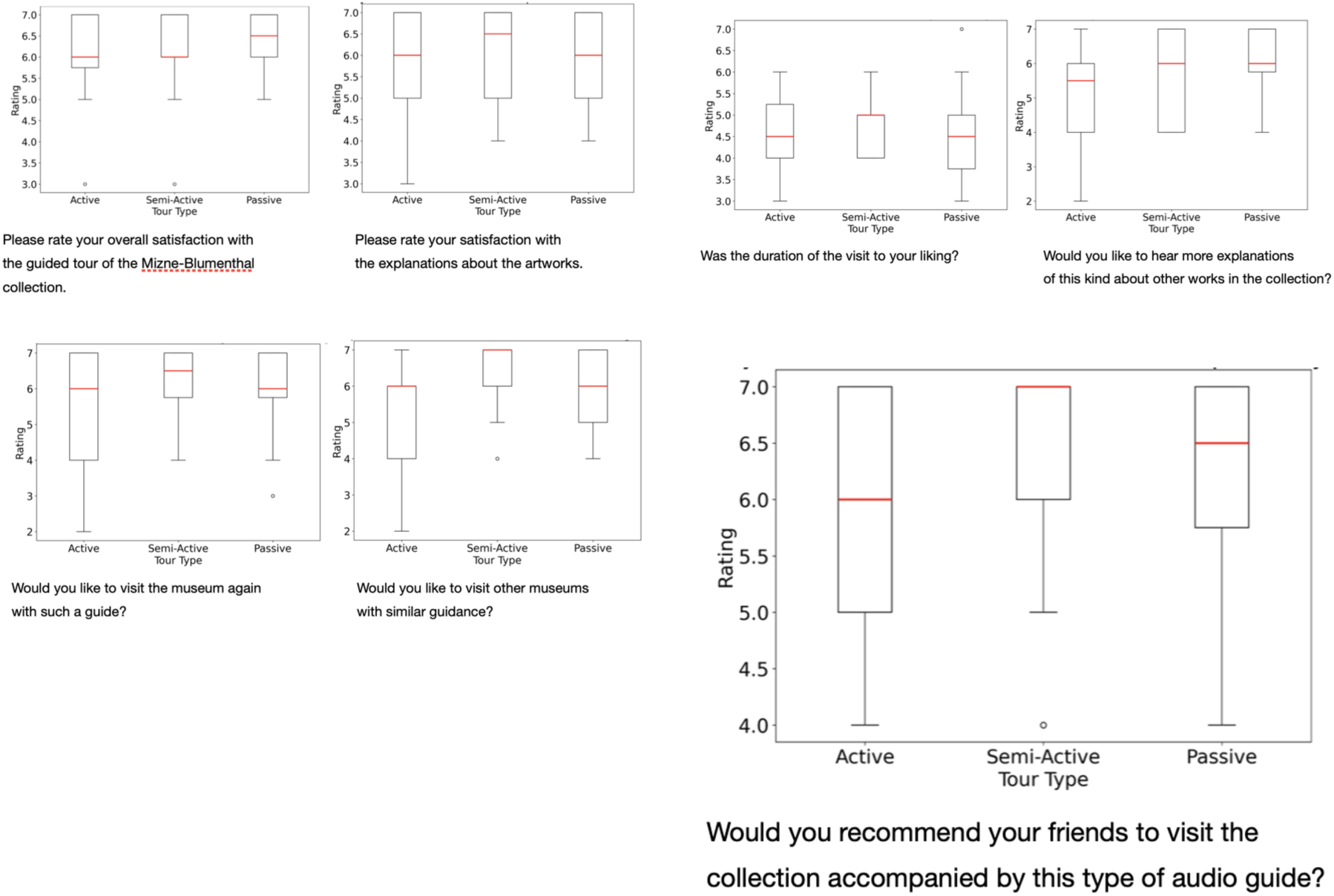
Post-tour questionnaire responses by tour type (Active, Semi-Active, Passive); medians in red. Only *“*willingness to visit other museums with similar guidance” differed reliably across conditions.

**Fig. 9.**
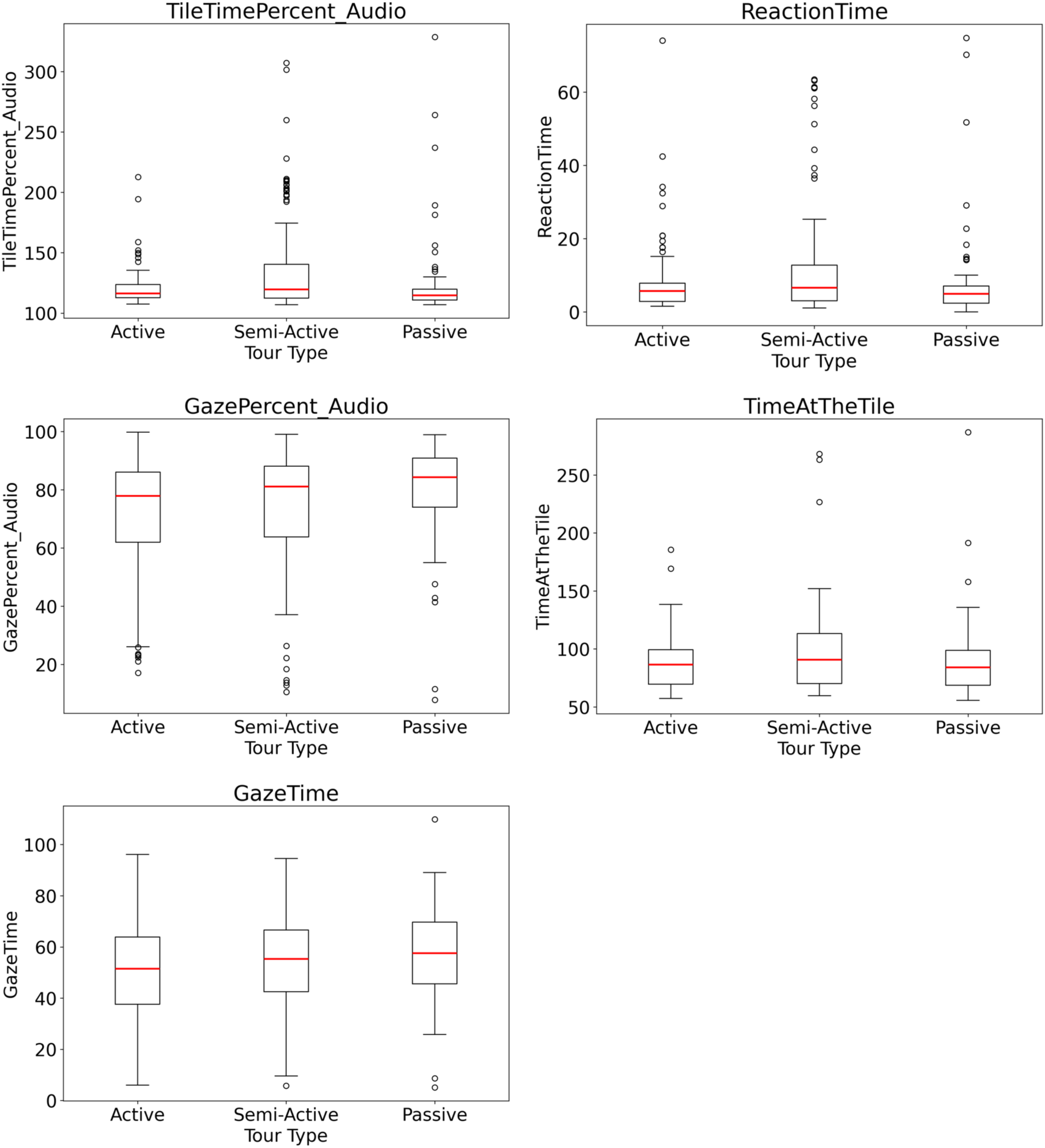
Painting-level viewing measures by tour type (medians in red): in-tile time as a percentage of audio duration, reaction time, gaze during the audio guide, time at the tile, and gaze time. Each differed reliably across conditions (all p ≤ 0.0225).

### 3.2 VR-derived measures and painting-level liking

We tested whether painting-level behavioral measures were associated with self-reported liking of the same painting (unit of analysis: the painting episode, N = 336). Because the route content was identical across conditions and liking distributions did not show reliable painting-by-painting shifts by tour type, we collapsed across tour type for the main painting-level analysis. Most VR-derived measures showed weak and inconsistent associations with liking (Table 3). Gaze during the audio guide showed a small positive trend (ρ = 0.106, p = 0.0533), whereas gaze time, reaction time, saccade rate, time at the tile, and in-tile time as a percentage of audio duration were not reliably related (all p ≥ 0.18).

**Table 3.** Spearman correlations between VR-derived measures and painting-level liking, collapsed across conditions (N = 336).

| Measure | Spearman $\rho$ | p |
| --- | --- | --- |
| GazePercent_Audio | 0.106 | 0.0533 |
| GazeTime | 0.072 | 0.1862 |
| ReactionTime | -0.052 | 0.3425 |
| SaccadeRate | -0.023 | 0.6719 |
| TimeAtTheTile | -0.015 | 0.7787 |
| TileTimePercent_Audio | 0.005 | 0.9206 |

Within-type analyses (N = 112 episodes per condition) revealed only two small, condition-specific effects: in the Semi-Active condition higher GazePercent_Audio was associated with higher liking (ρ = 0.218, p = 0.0208), and in the Passive condition shorter reaction time was associated with higher liking (ρ = −0.192, p = 0.043). These associations did not generalize to other measures or reappear in other conditions. Overall, gaze engagement during the audio guide was the clearest behavioral signal linked to liking, whereas movement-and timing-derived measures were weaker predictors of explicit preference.

We additionally analyzed the binary choice task, in which each tour painting was paired with an unseen work by the same artist; the mean of the binary outcome is interpretable as the proportion of participants who chose the tour painting (ranging from 0.13 for Braque to 0.75 for Picasso). Liking and binary preference were positively related for several paintings (e.g., de Chirico ρ = 0.62; Janco ρ = 0.43; Braque ρ = 0.41; all p < 0.01), indicating that the two response formats capture overlapping aspects of preference, though the strength varied by artwork. When predicting binary preference from behavior, self-reported liking was the strongest and most consistent correlate, and no single VR-derived measure emerged as a reliable predictor across paintings.

### 3.3 VR-derived measures and tour-level satisfaction

We next tested whether participant-level VR summaries related to overall tour satisfaction (one value per participant), computed as the average of the seven general post-tour items. Across the full sample (N = 48), no VR-derived measure showed a reliable relationship with satisfaction (Table 4); the strongest trends were negative associations with locomotion (time moving ρ = −0.242, p = 0.097; average speed ρ = −0.239, p = 0.103). Within-condition correlations (N = 16 each) were limited and inconsistent across conditions. Thus, *“*how people walked and looked” was not a robust proxy for how much they reported enjoying the visit.

**Table 4.** Spearman correlations between participant-level VR summaries and average general (tour-satisfaction) rating (N = 48).

| Measure | Spearman $\rho$ | p |
| --- | --- | --- |
| TimeMoving | -0.242 | 0.0973 |
| AvgSpeed | -0.239 | 0.1026 |
| TimeStill | 0.235 | 0.1085 |
| TotalDistance | -0.228 | 0.1198 |
| TotalExperimentTime | -0.156 | 0.2911 |
| AvgSaccadeRate | -0.139 | 0.3463 |
| AvgGazePercent_Audio | 0.130 | 0.3790 |
| AvgGazeTime | 0.059 | 0.6885 |

### 3.4 Effects of agency on self-report

Participants reported high satisfaction with the guided tour (M = 6.13) and the explanations (M = 5.96) and tended to endorse willingness to revisit (M = 5.76) and to recommend (M = 6.07); perceived duration was near the scale midpoint (M = 4.49). Comparing Active, Semi-Active, and Passive conditions on each questionnaire item (Kruskal–Wallis), no reliable differences were found for overall satisfaction, satisfaction with the explanations, perceived duration, willingness to revisit, or willingness to recommend (all p > 0.085). The only item differing across conditions was the broader attitude question *“*Would you like to visit other museums with similar guidance?” (H = 6.51, p = 0.0385). Painting-level liking ratings also did not differ reliably by tour type. Thus, agency did not reliably shift reported enjoyment of this tour or painting-level liking, though it may have influenced general openness to similar guided experiences beyond the current visit (Table 5).

**Table 5.** Kruskal–Wallis tests of tour-type effects (Active / Semi-Active / Passive). Top: selected self-report items. Bottom: painting-level viewing measures. Bold p-values are significant at α = .05.

| Outcome | H | p |
| --- | --- | --- |
| Overall tour satisfaction (self-report) | 1.08 | 0.5843 |
| Satisfaction with explanations | 0.46 | 0.7955 |
| Willingness to visit other museums w/ similar guidance | 6.51 | 0.0385* |
| Total experiment time (participant level) | 11.95 | 0.0025* |
| Total distance / avg speed / time moving (participant) | , | > 0.10 (n.s.) |
| TileTimePercent_Audio (painting level) | 16.47 | 0.0003* |
| ReactionTime (painting level) | 11.71 | 0.0029* |
| GazePercent_Audio (painting level) | 10.60 | 0.0050* |
| TimeAtTheTile (painting level) | 10.25 | 0.0059* |
| GazeTime (painting level) | 7.59 | 0.0225* |
| SaccadeRate (painting level) | 2.58 | 0.2755 |
| SelfReportedLiking (painting level) | 0.48 | 0.7882 |

### 3.5 Effects of agency on behavior

Finally, we tested whether agency shaped observable behavior inside VR. At the participant level, total experiment time differed across conditions (H = 11.95, p = 0.0025), whereas overall locomotion summaries (average speed, total distance, time still, time moving) did not (all p > 0.1). The time difference is best interpreted as a structural consequence of the manipulation: the Active and early Semi-Active conditions include time spent reading and answering choice questions, which could not be cleanly separated from *“*pure” viewing time. Within-type correlations between total experiment time and questionnaire outcomes were inconsistent across conditions, arguing against a simple *“*longer tours = more enjoyment” interpretation.

At the painting level, where the trial windows were constructed to exclude the choice screens, several time-and gaze-based measures differed reliably across conditions (Table 5): in-tile time as a percentage of audio duration (H = 16.47, p = 0.0003), reaction time to first fixation (H = 11.71, p = 0.0029), gaze during the audio guide (H = 10.60, p = 0.005), time at the tile (H = 10.25, p = 0.006), and gaze time (H = 7.59, p = 0.0225). In contrast, self-reported liking (p = 0.79) and saccade rate (p = 0.28) did not differ across conditions. Taken together, agency influenced exploratory and visual-engagement behavior during viewing episodes, where participants stood and how continuously they looked during the audio guide, without translating into a reliable increase in reported enjoyment.

## 4 Discussion

We tested whether VR-derived behavioral signals measured during a walkable, audio-guided museum tour map onto what participants later reported enjoying, and whether agency changed that enjoyment. Across the full dataset (N = 48; 336 painting segments), the clearest behavioral signal was gaze engagement during the audio guide, which showed a small positive trend with painting-level liking (GazePercent_Audio, p = 0.0533). Other VR-derived measures were weak and inconsistent predictors of explicit enjoyment outcomes, painting liking, preferring tour paintings over unseen ones, and overall tour satisfaction.

A key gap in prior work was that real-walking VR measurement had usually been demonstrated around a single artwork rather than embedded in a multi-artwork tour. Our trial-and-tile approach extends real-walk measurement to a full-visit structure: the tour unfolds across multiple paintings, and each painting yields a comparable analysis window defined by the intersection of its trial interval and the empirically defined tile in front of it. The pipeline produced sensible distributions at three levels, pooled painting segments, the same measures by artwork, and participant-level aggregates. Notably, in-tile time exceeding 100% of audio duration is not an artifact but reflects that the viewing episode often contains more than listening, including time before and after the narration spent standing in the tile.

The audio guide created a common temporal anchor: everyone heard the same narration per artwork, so gaze during that interval has a direct interpretation as visual engagement with the painting while being guided. One reading of the modest gaze–liking link is that keeping gaze on the canvas while listening may help bind verbal explanation to visual detail, supporting a more coherent and satisfying experience, though the effect was neither strong nor universal. The small overall size of the painting-level correlations marks an important boundary condition: explicit liking is shaped by many factors not captured by basic “where you stood” and “how much you looked” measures, such as prior taste, style preference, interpretation, and narrative resonance.

Locomotion metrics in an audio-guided tour are not pure indicators of interest. Walking may reflect navigation, comfort, habituation to VR, or simple repositioning, none of which necessarily implies greater liking; accordingly, total distance, average speed, and time moving were not reliable correlates of satisfaction. Even painting-proximal measures such as time in the tile are ambiguous, since participants may linger because they are attentive, because the narration is long, or because they are waiting for the next step. These results argue for treating movement and pacing as context variables rather than direct proxies for enjoyment, and for focusing on measures tied to the intended viewing moment (here, the audio-guide window).

### 4.1 The role of agency

We did not find reliable condition differences in tour satisfaction or painting-level liking, which contrasts with a broad literature showing that choice can enhance motivation and enjoyment [15–20]. Agency nonetheless clearly modulated viewing behavior: several painting-level time-and gaze-based measures differed across conditions (all p ≤ 0.0225). A crucial design detail is that the route itself was identical across conditions; only the presence of in-headset choice questions differed. This matters for interpreting total experiment time, which is expected to increase with the choice components in Active and Semi-Active and could not be cleanly decomposed. The main painting-level viewing measures are less vulnerable to this confound, because trials were constructed to exclude the choice questions while keeping the core viewing episode comparable across conditions.

### 4.2 Binary preference as a validity check

Alongside 1–7 liking ratings, we collected a binary choice preference measure to test whether a different response format yields a similar preference signal. Liking and binary preference were positively related for most paintings, supporting the idea that both capture overlapping aspects of subjective preference, although the strength varied by artwork. As with the continuous ratings, no single VR-derived measure emerged as a strong, consistent predictor of binary preference, reinforcing the conclusion that basic movement and gaze summaries only partially explain explicit preference.

### 4.3 Limitations and future directions

Several limitations define clear next steps. First, the between-subjects design yields relatively small per-condition samples, limiting sensitivity to subtle agency effects on enjoyment. Second, because the route was identical across conditions, the manipulation altered the experience of choosing rather than the content of the tour; future work could test stronger manipulations in which choices change the route outcome while still controlling for content exposure. Third, some locomotion measures are sensitive to tracking artifacts and occasional re-calibration; future studies could add quality control for locomotion summaries or focus on within-painting micro-movements that may better reflect viewing strategy than total distance. Finally, the facial-expression channel logged here was not analyzed; because all streams are time-aligned, they can be segmented with the same trial-and-tile windows and integrated into multimodal models. A key direction is to extend the present pipeline with richer gaze features and affective channels to test whether enjoyment and preference become more predictable when behavioral and affective signals are combined.

## 5 Conclusion

This study asked whether VR-derived behavioral signals are linked to enjoyment during a virtual museum tour, and whether increasing agency changes that enjoyment. At the participant level, aggregated VR measures showed no reliable association with tour satisfaction; at the painting level, gaze engagement during the audio guide showed a small positive trend with liking (p = 0.0533). Agency did not reliably increase self-reported enjoyment of the tour or painting-level liking, but it did alter behavior: several painting-level viewing measures differed across conditions (all p ≤ 0.0225). Together these results suggest a dissociation between subjective enjoyment and the micro-structure of viewing in VR, choice can change pacing and visual engagement without changing how much people report liking the experience. The main contribution is methodological: real-walking VR can support a full-tour, painting-level analysis framework that preserves ecological behavior while enabling structured, artwork-specific measurement, providing a foundation for richer models of attention, affect, and enjoyment in future VR museum research.

## Declarations

Ethics approval The study was approved by the Tel Aviv University ethics committee. All participants provided written informed consent prior to participation.

Data and code availability Processed data tables and the full analysis code are available at github.com/Slab Museum/Enjoyment. Logging templates are available at minervaxr.sites.tau.ac.il/open-science.

## Author contributions

Y.S. and T.S. conceived and designed the study. J.H. contributed to the conception of the project and recorded the audio-guide narration. Y.S. collected the data, implemented the analysis pipeline, and performed the analyses. Y.S. and T.S. interpreted the results and drafted the manuscript. T.S. supervised the project and acquired funding. All authors revised and approved the final manuscript.

## Acknowledgements

We would like to deeply thank Tania Coen-Uzzielli, Director of the Tel Aviv Museum of Art, for early discussions on the topic, ideation and for providing digital materials. We would like to thank Noa Barel for developing and maintaining the Unity application and for technical support throughout.

